# Cdc48 and its co-factor Ufd1 segregate CENP-A from centromeric chromatin and can induce chromosome elimination in the fission yeast *Schizosaccharomyces pombe*

**DOI:** 10.1101/2023.06.15.545183

**Authors:** Yukiko Nakase, Hiroaki Murakami, Michiko Suma, Kaho Nagano, Airi Wakuda, Teppei Kitagawa, Tomohiro Matsumoto

## Abstract

CENP-A, a variant of histone H3, determines the identity of the centromere, a chromosome locus at which a microtubule attachment site termed kinetochore is assembled. Because the position and size of the centromere and its number per chromosome must be maintained for the faithful segregation of chromosomes, the distribution of CENP-A is strictly regulated. In this study, we have aimed to understand mechanisms to regulate the distribution of CENP-A by a genetic approach in the fission yeast *Schizosaccharomyces pombe*. A mutant of the *ufd1^+^* gene (*ufd1-73*) encoding a cofactor of Cdc48 ATPase is sensitive to CENP-A expressed at a high level and allows mislocalization of CENP-A. ChIP analysis has revealed that the level of CENP-A in centromeric chromatin is increased in the *ufd1-73* mutant even when CENP-A is expressed at a normal level. A preexisting mutant of the *cdc48^+^* gene (*cdc48-353*) phenocopies the *ufd1-73* mutant. We have also shown that Cdc48 and Ufd1 proteins physically interact with centromeric chromatin. Finally, Cdc48 ATPase with Ufd1 artificially recruited to the centromere of a mini-chromosome (*Ch16*) induce a loss of CENP-A from *Ch16*, resulting in an increased rate of chromosome loss. It appears that Cdc48 ATPase, together with its cofactor Ufd1 segregates excess CENP-A from chromatin, likely in a direct manner, to maintain proper distribution of CENP-A. This mechanism may play an important role in centromere disassembly, a process to eliminate CENP-A massively to inactivate the kinetochore function during development, differentiation, and stress response in other organisms.

**Significance statement:** Maintaining the proper distribution of CENP-A is crucial for centromere identity. This process involves accurate positioning of CENP-A and removal of excess CENP-A from chromatin. The Cdc48-Ufd1 complex is essential for this regulation as it acts as a segregase that directly eliminates surplus CENP-A from chromatin. These findings have therapeutic significance as targeting the Cdc48 complex can potentially correct abnormal karyotypes in disomic embryos and prevent trisomic disorders like Down syndrome. This research advances our understanding of centromeric chromatin regulation and its potential therapeutic applications.

## Introduction

The centromere is a specialized chromosomal locus at which a microtubule attachment site termed kinetochore is assembled in mitosis. CENP-A is a variant of histone H3 preferentially found in centromeric chromatin and serves as a base of the kinetochore assembly (1–3). Because CENP-A determines the identity of the centromere, its distribution must be strictly regulated. The position and size of the centromere and its number per chromosome must be maintained for the faithful segregation of chromosomes. CENP-A distribution is likely regulated by two mechanisms, one to deposit CENP-A at the correct positions and another to remove excess CENP-A from chromatin.

Interestingly, accumulating evidence suggests that CENP-A is sometimes massively eliminated from the centromeric chromatin. In mouse NIH/3T3 cells treated with genotoxic reagents, CENP-A is eliminated from the centromeric chromatin and found at the periphery and inside the nucleolus. This process requires the ATM-mediated signaling pathway and chromatin remodeling factors. CENP-A elimination is considered a part of DNA damage response, which safeguards the genome (4). In a holocentric organism, *Ascaris suum*, a part of genomic DNA is eliminated in somatic precursor cells during early development. At regions that will be lost, CENP-A is dramatically diminished, resulting in the absence of kinetochores and nondisjunction of these regions in mitosis (5). Elimination of CENP-A also plays a vital role in the differentiation of *Arabidopsis thaliana*. Non-dividing pollen vegetative cells undergo loss of CENP-A dependently on the activity of Cdc48A proteins. The disappearance of CENP-A from the centromeres is accompanied by the decondensation of centromeric heterochromatin and activation of rRNA genes. Elimination of CENP-A is proposed to contribute to both the growth and prevention of cell division of the pollen cells (6).

Cdc48, also known as p97 or valosin-containing protein (VCP), is a member of the ATPase associated with diverse cellular activities (AAA) ATPase family (7, 8). It utilizes intrinsic ATPase activity and mediates the extraction of proteins from cellular compartments or protein complexes for recycling or degradation. Although Cdc48 ATPase is best known for its function in endoplasmic reticulum-associated degradation (ERAD), it also plays roles at chromatin (9).

We have previously reported that the Rpt3 protein, a subunit of the 19S proteasome, is involved in regulating the distribution of CENP-A (10). The *rpt3-1* mutant was identified through a screen for mutants that became temperature sensitive upon overexpression of CENP-A. ChIP analysis revealed that CENP-A was distributed broader and denser in the centromeric chromatin of the *rpt3-1* mutant, even when CENP-A was expressed at a normal level.

In this study, we report that Cdc48 ATPase and its cofactor Ufd1 are required to maintain proper distribution of CENP-A. In a mutant of the *ufd1^+^* gene, isolated from the screen the same as the *rpt3-1* mutant, the level of CENP-A in centromeric chromatin is increased. Consistently, a mutant in the *cdc48*^+^ gene (*cdc48-353*) exhibits similar phenotypes. We have also found that both Cdc48 and Ufd1 proteins associate with the centromeric regions as well as other chromosomal regions. Finally, Cdc48 ATPase with Ufd1 artificially recruited to the centromere of a mini-chromosome (Ch16) induce a loss of CENP-A from Ch16, resulting in an increased rate of chromosome loss. We propose that Cdc48 ATPase and its cofactor segregate excess CENP-A from chromatin, likely directly, to maintain proper distribution of CENP-A.

## Results

### Centromeric chromatin in the *ufd1-73* and *cdc48-353* mutants

Our preliminary examination suggested that the *ufd1-73* and *cdc48-353* mutants could not remove excess Cnp1 from chromatin (Fig. S1). Because we attempted to investigate the role of Ufd1 and Cdc48 in the maintenance of proper distribution of Cnp1 in the centromeric chromatin under physiological conditions, the experiments described below were performed with cells expressing Cnp1 tagged with GFP from the native promoter. When Cnp1 was solely expressed from its native promoter, the *ufd1-73* and *cdc48-353* grew slightly slower than the wild-type strain at 32 °C but could not grow at 36 °C (Fig. 1A).

**Figure 1.**
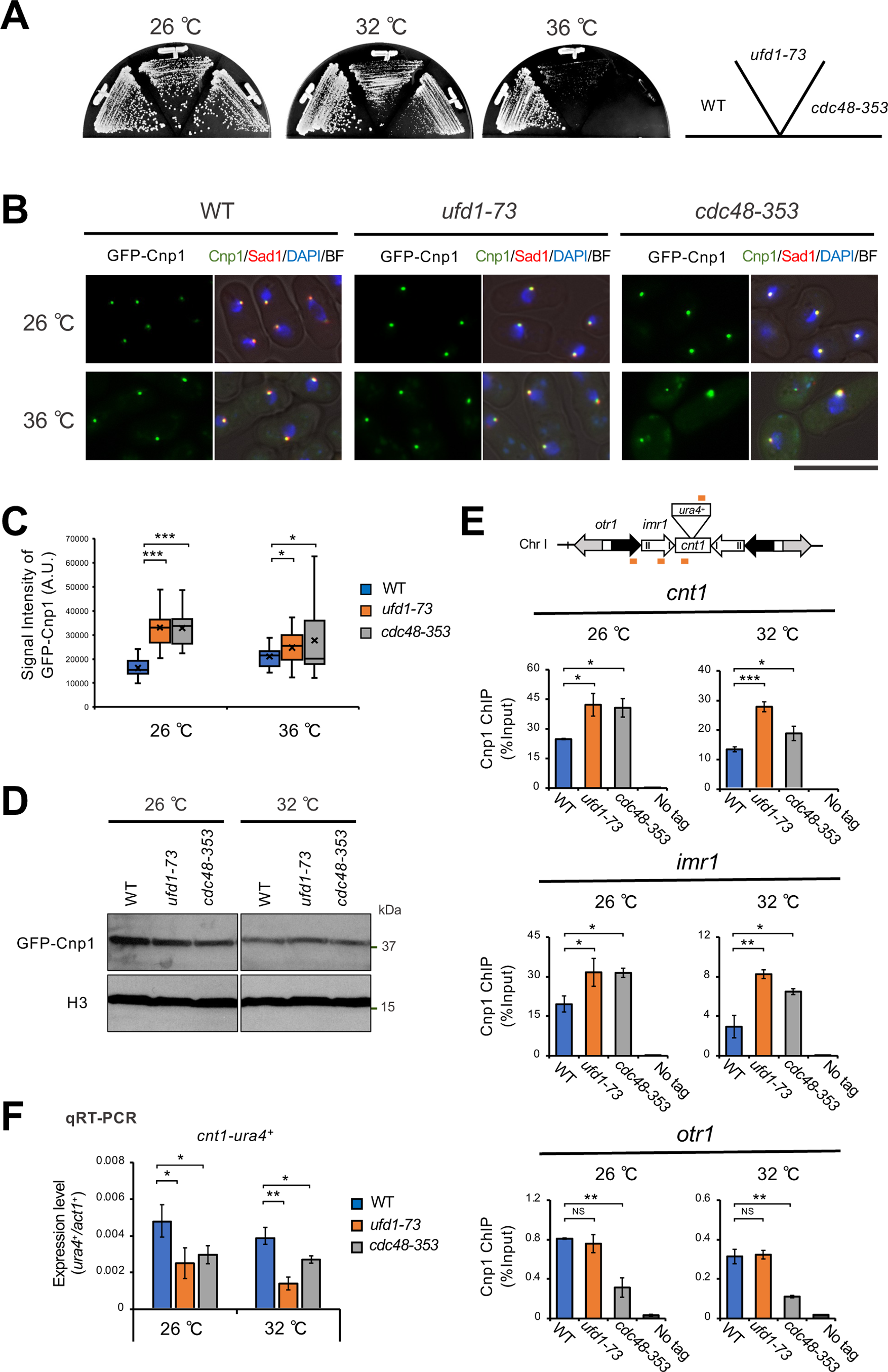
Distribution of Cnp1 in *ufd1-73* and c*dc48-353*. (A) Temperature sensitivity of *ufd1-73* and *cdc48-353* mutants. Strains were streaked on YEA plates. (B) Localization of GFP-Cnp1 from a native promoter. Strains expressing GFP-Cnp1 from native promoter were grown to mid-log phase in liquid YE medium at 26 °C and shifted to 36 °C for 6 hr. Sad1-mCherry: SPB maker. Scale bar: 10 μm. (C) The statistic analysis of (B). The GFP-Cnp1 intensity was calculated using NIH ImageJ and shown in arbitrary units. Representative experiments are shown (n = 3). (D) The protein level of GFP-Cnp1 by western blot. Strains were grown to mid-log phase in liquid YE medium at 26 °C and shifted to 32 °C for 8 hr. (E) Localization of GFP-Cnp1 by ChIP analysis. Strains were grown to mid-log phase in liquid YE medium at 26 °C and shifted to 32 °C for 3 hr. All data represent the mean±s.e.m. (n = 3). The orange bars indicate the position of primers used for ChIP or qRT-PCR (F). (F) Gene silencing of centromeric regions. Strains were grown as (E). qRT-PCR analyzed mRNA levels. Data are presented as mean±s.e.m. (n = 3). *P*-values (unpaired *t*-test) comparing controls (WT): \**p* < 0.05, \*\**p* < 0.01, \*\*\**p* < 0.001, NS: not significant.

Microscopic observation of the mutants revealed that the signal from GFP-Cnp1 appeared as a single speckle but that the intensity of the fluorescent signal was about twice stronger as that of the wild-type cells at both 26 °C and 36 °C (Fig. 1B & C). We also examined the level of GFP-Cnp1 by immunoblot with an antibody to GFP. As shown in Fig. 1D, the level of GFP-Cnp1 in the mutants was comparable to that of the wild-type cells.

In fission yeast, each centromere contains a central domain (*cnt*) flanked by innermost repeats (*imr)*, which provide the core of kinetochore formation. This core domain is surrounded by arrays of outer repeats (*otr*). Previous studies showed that Cnp1 is preferentially incorporated into *cnt* and *imr* but poorly into *otr*. ChIP analysis revealed that the amount of Cnp1 in *cnt* and *imr* chromatin of the mutants was increased (Fig. 1E). Gene silencing at the central domain (*cnt*) correlates with the amount of Cnp1 incorporated into the *cnt*-chromatin (4). As shown in Fig. 1F, the expression of the *ura4^+^*gene inserted at the *cnt* domain of *cen1* was silenced at 26 °C and 32 °C in the mutants.

The results described above indicated that while the *ufd1-73* and *cdc48-353* mutants expressed Cnp1 at a level similar to that of the wild-type strain, their centromeric chromatin at *cnt* and *imr* domains excessively incorporated Cnp1. Cdc48 ATPase and its cofactor, Ufd1, might be required to maintain proper distribution of Cnp1 at the centromere.

### Localization of Ufd1 and Cdc48

A previous study demonstrated that the Cdc48 proteins were localized at chromatin and in the cytoplasm in fission yeast (11). We reinvestigated the localization of Cdc48 with Ufd1 and found that both GFP-tagged Cdc48 protein and RFP-tagged Ufd1 protein were localized in the chromatin domain as well as the cytoplasm (Fig. 2A). However, microscopic observation could not provide clear evidence for localization of the two proteins at centromeric chromatin. We, therefore, examined the association of Cdc48 and Ufd1 with centromeric chromatin more precisely by ChIP analysis. As shown in Fig. 2B, Cdc48 and Ufd1 proteins were found to localize to the centromeric regions as well as other chromosomal regions, *18S,* subtelomere, and *act1^+^*.

**Figure 2.**
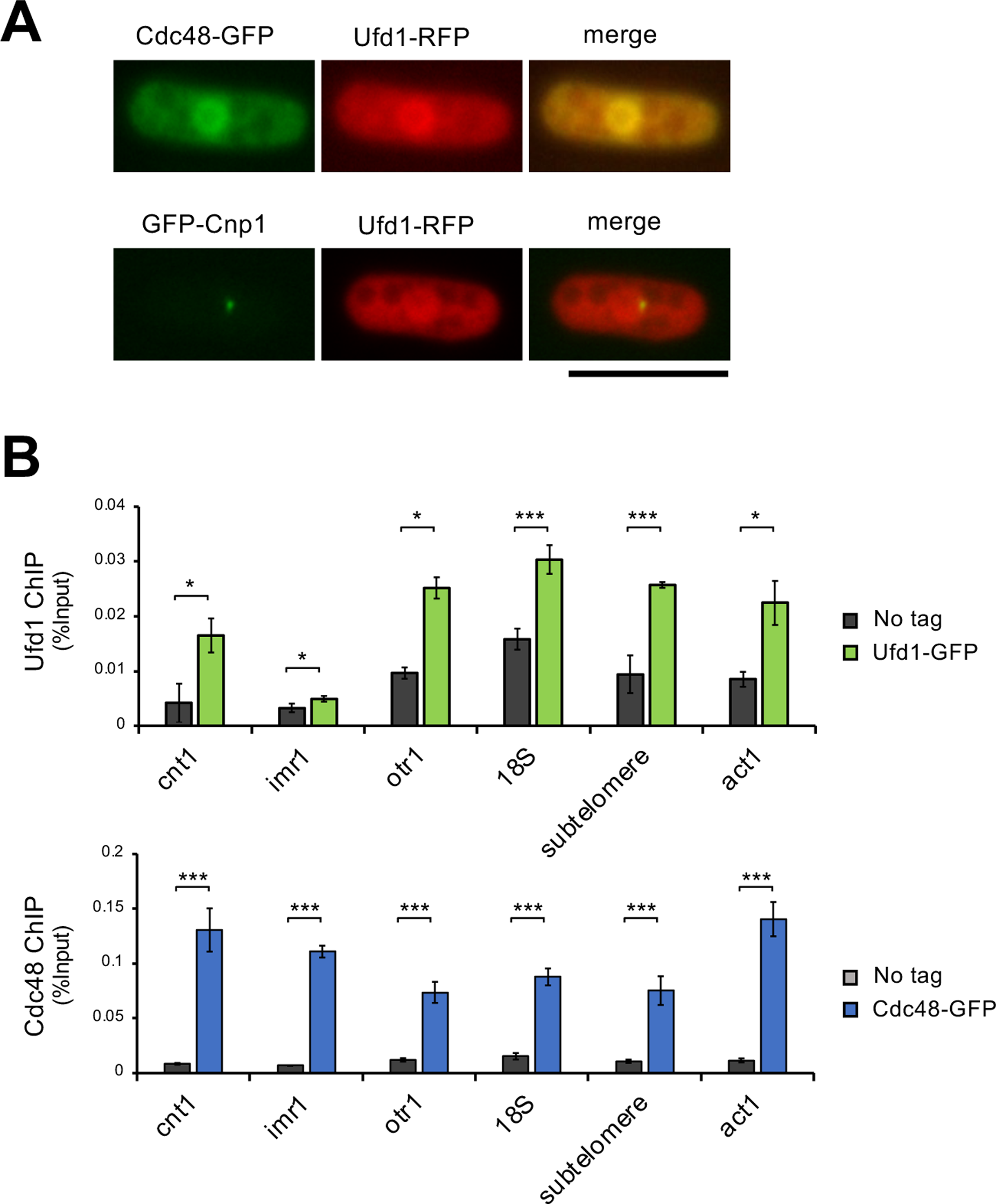
Localization of Ufd1 and Cdc48. (A) Localization of Ufd1-RFP, Cdc48-GFP, and GFP-Cnp1. Strains were grown to mid-log phase in liquid YE medium. Scale bar: 10 μm. (B) Localization of Ufd1-GFP and Cdc48-GFP by ChIP analysis. Strains were grown to mid-log phase in liquid YE medium at 26 °C. All data represent the mean±s.e.m. (n = 3). A two-sided unpaired *t*-test was used to calculate all *P*-values. \**p* < 0.05, \*\**p* < 0.01, \*\*\**p* < 0.001, NS: not significant.

### Removal of Cnp1/CENP-A by Cdc48 segregase

Cdc48 ATPase with Ufd1 is a segregase that extracts proteins from cellular compartments or protein complexes. Because our results described above indicated that Cdc48 ATPase and Ufd1 are associated with chromatin, and their dysfunction allows excess accumulation of Cnp1, we hypothesized that Cdc48 ATPase with Ufd1 segregates CENP-A from chromatin. To test this hypothesis, we attempted to increase the level of Cdc48 segregase at centromeric chromatin and examine the amount of Cnp1. We ectopically expressed cnp3c-GFP-Ufd1, a fusion protein consisting of the C-terminal domain (414-643 aa) of Cnp3 previously shown to have a centromere binding activity (12), GFP as a visible marker, and Ufd1. As shown in Fig. 3A, the cnp3c-GFP-Ufd1 fusion protein was localized throughout the chromatin domain and concentrated at the centromeres. In separate experiments, we examined the cnp3c-Ufd1 fusion protein for its ability to recruit Cdc48 ATPase and Npl4, another cofactor functioning with Ufd1. As shown in Fig. 3B, Cdc48 ATPase tagged with GFP was concentrated at the centromeres when the cnp3c-Ufd1 fusion protein was ectopically expressed. Likewise, Npl4 tagged with mCherry was focused at the centromeres (Fig. 3C). Together, these results suggested that the three components (Cdc48, Ufd1, and Npl4) required for the functional Cdc48 segregase were recruited to centromeric chromatin by ectopic expression of the cnp3c-Ufd1 fusion protein. We next examined the three components for their activity to segregate Cnp1. As shown in Fig. 3D and E, the fluorescent signal from GFP-Cnp1 in cells expressing the cnp3c-Ufd1 fusion protein was less than half that of cells expressing cnp3c alone. Although the expression level of cnp1 was not changed, the reduction of Cnp1 in centromeric chromatin was confirmed by ChIP (Fig. 3G & H). Due to the decrease of Cnp1 at the centromeres, approximately 10 % of the cells expressing the cnp3c-Ufd1 fusion protein exhibited abnormal chromosome segregation (Fig. 3F). Furthermore, gene silencing of the *ura4^+^* gene inserted at *cnt1* was loosened under expression of cnp3c-Ufd1 (Fig. 3I). These results consistently indicated that the three components recruited to the centromeres by ectopic expression of cnp3c-Ufd1 could function as a segregase to extract CENP-A from chromatin.

**Figure 3.**
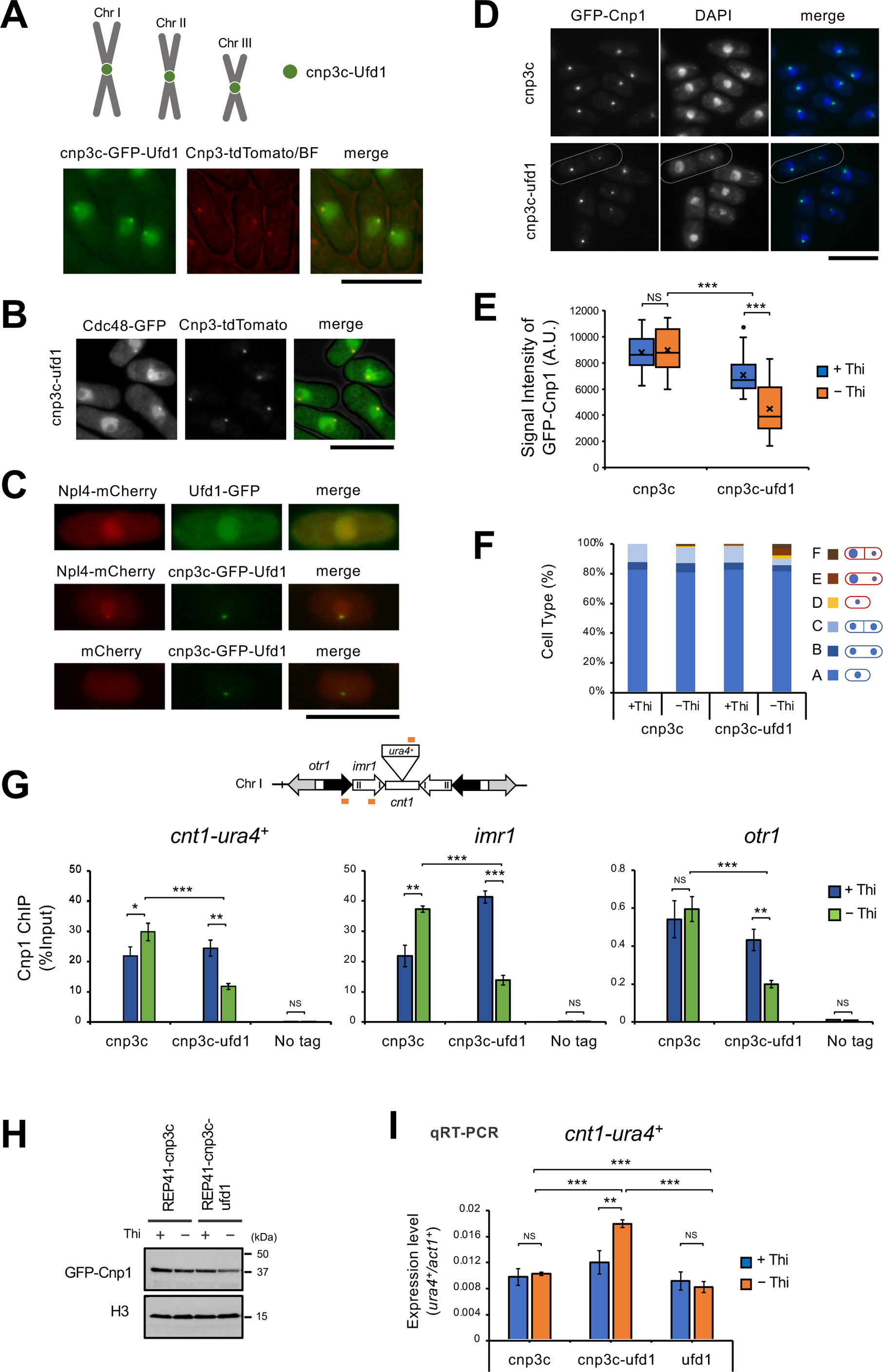
Ufd1 inhibits the chromatin binding of Cnp1. (A) Localization of cnp3c-GFP-Ufd1. The C-terminal domain (414-643 aa) of Cnp3 indicates centromere binding activity (12). We constructed the C-terminal domain of Cnp3 fused *ufd1^+^* plasmids. WT strains were transformed with pREP81-cnp3c-GFP-Ufd1. The resulting transformants were grown to mid-log phase in liquid EMM+thiamine medium at 26 °C and then transferred to EMM-thiamine medium for induction for 24 hr. (A-I) was observed under the same conditions. (A-D) scale bars: 10 μm. (B) Localization of Cdc48-GFP under expressing cnp3c-Ufd1. (C) Localization of Npl4-mCherry under expressing cnp3c-GFP-Ufd1. (D) Localization of GFP-Cnp1 under expressing cnp3c-Ufd1. White dotted lines indicate unevenly distributed cells. (E) The statistic analysis of the signal intensity of GFP-Cnp1 in (D). (F) Cell types in (D). Cell type A-C: normal cells; D-F: abnormal cells. The GFP-Cnp1 intensity was calculated using NIH ImageJ and shown in arbitrary units. Representative experiments are shown (n = 3). (G) Localization of GFP-Cnp1 by ChIP analysis under expressing cnp3c-Ufd1. All data represent the mean±s.e.m. (n = 3). The orange bars indicate the position of primers used for ChIP. (H) The protein level of GFP-Cnp1 by western blot under expressing cnp3c-Ufd1. (I) Gene silencing of centromeric regions under expressing cnp3c-Ufd1. qRT-PCR analyzed mRNA levels. Data are presented as mean±s.e.m. (n = 3). A two-sided unpaired *t*-test was used to calculate all *P*-values. \**p* < 0.05, \*\**p* < 0.01, \*\*\**p* < 0.001, NS: not significant.

### Elimination of a mini chromosome by Cdc48 segregase

We finally assessed the ability of the Cdc48 segregase to eliminate a specific chromosome by extracting Cnp1/CENP-A. Fission yeast mini-chromosome *ch16* is a derivative of chromosome III, which lacks most of the arm regions but maintains the centromere III (13). It is a model chromosome transmitted stably and is not essential for viability.

A plasmid expressing tetR-Ufd1-mcherry, a fusion protein consisting of tetR, Ufd1, and mCherry as a visible marker, was transformed into a strain in which *tetO* array with *ura4^+^* cassette was inserted at the *imr3L* domain of *ch16.* We anticipated that Ufd1 could be recruited to the *imr3L* domain by tetR, which recognized *tetO* arrays. As shown in Fig. 4A, upon induction of the fusion protein, tetR-Ufd1-mCherry, the fluorescent signal from mCherry appeared as a single speckle and was colocalized with GFP-Cnp1, suggesting that Ufd1 was recruited to the centromere of *ch16* as anticipated. The tetR-Ufd1-mCherry fusion protein could also recruit Cdc48-GFP to the centromeric region of *ch16* (Fig. 4B and C). Based on these results, we speculated that the functional Cdc48 segregase was recruited to the centromere of *ch16*. We then examined the level of Cnp1 at the centromere of *ch16*. As shown in Fig. 4D and E, induction of the tetR-Ufd1-mCherry fusion protein significantly decreased the level of Cnp1 at the centromere of *ch16,* but did not affect the centromere of Chromosome I. The reduction in the level of Cnp1 affected the function of the centromere of *ch16*. We estimated the loss rate of *ch16* per division was approximately ten-fold higher than that of cells expressing tetR-mCherry, a fusion protein lacking Ufd1 (Fig.4E and F). These results suggested that the Cdc48 segregase disassembled the centromeric chromatin and induced the elimination of *ch16*.

**Figure 4.**
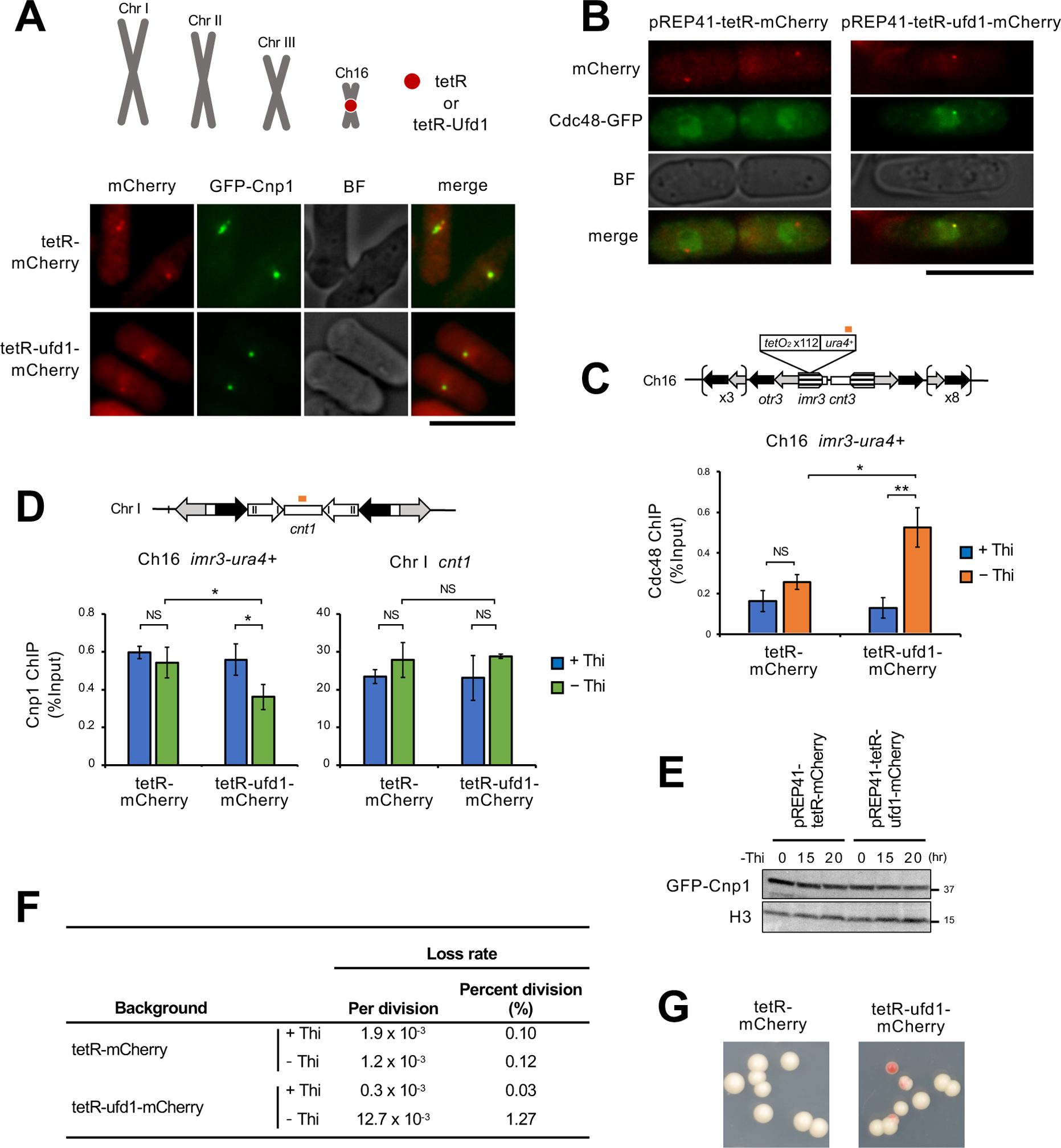
Artificial enrichment of Ufd1 at the centromere causes chromosome instability. (A) Artificial enrichment of Ufd1 at the centromere of mini-chromosome *ch16* using tetO/tetR system. We constructed tetR fused *ufd1^+^* plasmids. The strains containing *tetO* sequence inserted centromeric region were transformed with pREP41-tetR-mCherry or pREP41-tetR-Ufd1-mCherry. The resulting transformants were grown to mid-log phase in liquid EMM+thiamine medium at 30 °C and then transferred to EMM-thiamine medium for induction for 24 hr. (A-D) was observed under the same conditions. Scale bars: 10 μm. (B) Localization of Cdc48-GFP under expressing tetR-Ufd1-mCherry. (C) Localization of Cdc48-GFP by ChIP analysis under expressing tetR-Ufd1-mCherry. All data represent the mean±s.e.m. (n = 3). The orange bar indicates the position of the primer used for ChIP. (D) Localization of GFP-Cnp1 by ChIP analysis under expressing tetR-Ufd1-mCherry. All data represent the mean±s.e.m. (n = 3). The orange bar indicates the position of the *cnt1* primer used for ChIP. The primer of *imr3-ura4^+^* used for ChIP was the same as (C). (E) The protein level of GFP-Cnp1 by western blot under expression terR-Ufd1-mCherry. The expression level of Cnp1 was not changed. Ufd1-cherry does not affect the expression level of Cnp1. (F) Stability of mini-chromosome. The stability of *ch16* was determined on YE-Ade media at 30 °C. (G) The colonies on YE-Ade under expressing tetR-mCherry or tetR-Ufd1-mCherry. A two-sided unpaired *t*-test was used to calculate all *P*-values. \**p* < 0.05, \*\**p* < 0.01, NS: not significant.

## Discussion

In this study, Cdc48 and its cofactor Ufd1 have been characterized with a focus on their role at the centromeric chromatin. We have shown by performing ChIP that a subset of each of the two proteins is localized at the centromeric chromatin. In the corresponding mutants (*cdc48-353* and *ufd1-73*), CENP-A (fission yeast Cnp1) excessively accumulates in the centromeric chromatin at the *cnt* and *imr* but not at the *otr* domains. These two lines of evidence suggest that Cdc48, along with Ufd1, play an important role in the maintenance of proper distribution of CENP-A at the core of the centromeric chromatin. We have previously reported that the 26S proteasome (or 19S proteasome subcomplex) associates with the centromeric chromatin and regulates the distribution of CENP-A (10). Interestingly, the *otr* domain of a mutant in the gene encoding Rpt3, a subunit of the 19S subcomplex, is more prone to excessive accumulation of CENP-A than the other two domains, *cnt,* and *imr*. These observations suggest that the level of CENP-A at the *otr* domain is under the surveillance of the 26S proteasome (or 19S proteasome subcomplex) but not of Cdc48 with Ufd1.

It has been reported that the Cdc48-Ufd1-Nlp4 complex removes Aurora B (14–16), CDT1 (17, 18), and DNA damage-associated factors (19, 20) from chromatin. We propose that the Cdc48 complex targets Cnp1/CENP-A as well and removes excess CENP-A from chromatin. It should be noted that Cdc48c, an isoform of Cdc48 in *Arabidopsis thaliana*, is required to eliminate CENP-A in non-dividing pollen vegetative cells (6). The function of Cdc48 with its co-factors as the CENP-A segregase is likely conserved through evolution.

To demonstrate more directly that the Cdc48 complex serves as the CENP-A segregase, we recruited the Cdc48 complex to the centromeric chromatin and monitored the level of Cnp1/CENP-A. When the Cdc48 complex was recruited to the centromeric chromatin of the regular chromosomes, the level of CENP-A at the *cnt* domain of chromosome I decreased to approximately 50 %. Likely due to the loss of CENP-A from the centromeric chromatin, some chromosomes lost the centromeric activity and segregated abnormally. Likewise, when the Cdc48 complex was recruited specifically to an artificial chromosome, *ch16*, the level of CENP-A at the *imr* domain of *ch16* decreased by approximately 30 %, and the loss rate of *ch16* increased by 10-fold. These results suggest that the Cdc48 complex is a CENP-A segregase.

A disomic embryo followed by fertilization with a normal sperm is a cause of disomic diseases, such as Down syndrome, Edward syndrome, and Patau syndrome. The Cdc48 complex, if targeted to the centromere of one of the disomic chromosomes in the embryo, may eliminate it, just like *ch16* shown in this study, and correct the karyotype of the embryo. This study may thus provide a basis for an approach to prevent trisomic diseases.

## Acknowledgments

We thank S Oda for the technical assistance, T Yuasa, and members of the Matsumoto lab for helpful discussion. We also thank NBRP Japan for the strains.

## Competing Interests

The authors declare no competing interests.

## Author Contributions

Y.N. and T.M. designed the research; Y.N., H.M., M.S., K.N., A.W., and T.K. performed the research; and Y.N. and T.M. wrote the paper.

## Funding

This work was supported by a grant from the Japan Society for the Promotion of Science (20K06485) to T. M. and Y.N.

## Material and Methods

### Yeast strains and growth media

The *S. pombe* strains used in this study are listed in Supplementary Table 1. As described previously, *S. pombe* cells were grown in YEA and EMM containing the appropriate nutrient supplements (21). All yeast transformations were performed with the lithium acetate method (22, 23).

### Plasmid construction

Plasmid pREP81-cnp3c-GFP-ufd1 was constructed as follows. The Gibson assembly inserted the ufd1 cDNA (−1 kb) into the plasmid which the C-terminus 414-643 aa of *cnp3* (0.8 kb) and GFP (0.7 kb) are linked downstream of the nmt81 promoter. Plasmid pREP41-cnp3c-ufd1 was constructed as follows. The Gibson assembly inserted the ufd1 cDNA (−1 kb) into the plasmid, which the C-terminus 414-643 aa of *cnp3* are linked downstream of the nmt41 promoter.

Plasmid pREP41-tetR-ufd1-mCherry was constructed as follows. A 0.6 kb *Sal*I(*BamH*I)-*Not*I fragment containing tetR was cloned into the corresponding site of pREP41-mCherry. A 1 kb *BamH*I-*Not*I fragment of *ufd1* cDNA was cloned into the corresponding site of pREP41-tetR-mCherry.

### Isolation of the *ufd1-73* mutant

Isolation of the *ufd1-73* was performed as described previously (24).

### Isolation of the *ufd1^+^* gene

A genomic DNA fragment that complements the temperature sensitivity was isolated from a fission yeast genomic DNA library using temperature sensitivity as a selection marker. The integration mapping proved that this fragment originated from the *ufd1-73* locus. A PCR-based strategy was used to identify the mutation site within the *ufd1^+^* coding sequence. The *ufd1-73* mutant gene contained a mutation of G55.

### Western blotting

As described previously, crude cell extracts were prepared from *S. pombe* (25). Polypeptides were resolved by SDS-PAGE gel and then transferred onto nitrocellulose membranes. Antibodies for western blotting were diluted as follows: mouse anti-GFP (Roche) 1:1000; rabbit anti-H3 (1729, abcam) 1:1000. Blots were developed using ECL reagents (Thermo).

### Microscopy

Images were acquired on a LEICA DM5500B (Leica) microscope with a HAMAMATSU ORCA-ER camera and a KEYENCE (BZ-8000).

### ChIP assay

ChIP was performed as described previously (10). The nucleotide sequences of the primer sets used in this study are listed in Supplementary Table 2.

### qRT-PCR

qRT-PCR was performed as described previously (26). The *ura4* primer sets are located in the region (0.2 kb) lacking in *ura4-DS/E*.

### Mini-chromosome stability assay

Mini-chromosome stability assay was performed as described previously (24).

## Notes

### Competing Interest Statement

The authors have declared no competing interest.

