## Supplemental text_YN for "Cdc48 and its co-factor Ufd1 segregate CENP-A from centromeric chromatin and can induce chromosome elimination in the fission yeast *Schizosaccharomyces pombe*": supText(ufd1)_YN0615_2023.docx

**Supplementary Materials and Methods**

Nakase et al., 2023

**Yeast strains and growth media**

The *S. pombe* strains used in this study are listed in Supplementary Table 1. *S. pombe* cells were grown in YEA and EMM containing the appropriate nutrient supplements as described previously (21). All yeast transformations were performed with the lithium acetate method (22, 23).

**Western blotting**

Crude cell extracts were prepared from *S. pombe* as described previously (24). Polypeptides were resolved by SDS-PAGE gel and then transferred onto nitrocellulose membranes. Antibodies for western blotting were diluted as follows: mouse anti-GFP (Roche) 1:1000; rabbit anti-H3 (1729, abcam) 1:1000. Blots were developed using ECL reagents (Thermo).

**Microscopy**

Images were acquired on a LEICA DM5500B (Leica) microscope equipped with a HAMAMATSU ORCA-ER camera and a KEYENCE (BZ-8000).

**ChIP assay**

ChIP was performed as described previously (10). The nucleotide sequences of the primer sets used in this study are listed in Supplementary Table 2.

**Supplementary Table 1. *S. pombe* strains used in this study**

| **Strain** | **Genotype** | **Source** |
| --- | --- | --- |
| **SP6** | *h^-^ leu1-32* | Matsumoto *et al*., (2002) (27) |
| **TK28** | *h^-^ lys1::nmt1-GFP-cnp1-lys1^+^ leu1-32* | Kitagawa *et al*., (2014) |
| **AW1** | *h^-^ lys1::nmt1-GFP-cnp1-lys1^+^ leu1-32 ufd1-73* | This study |
| **YKK1114** | *h^-^ lys1::nmt1-GFP-cnp1-lys1^+^ leu1-32 cdc48-353* | This study |
| **YKK1143** | *h^-^ leu1::nmt1-GFP-h3-leu1^+^* | This study |
| **YKK1142** | *h^-^ leu1::nmt1-GFP-h3-leu1^+^ ufd1-73* | This study |
| **YKK1151** | *h^-^ leu1::nmt1-GFP-h3-leu1^+^ cdc48-353* | This study |
| **MS122** | *h^-^ lys1::nmt1-GFP-cnp1-lys1^+^ sad1-mCherry::kan^R^ leu1-32* | This study |
| **YKK1126** | *h^-^ lys1::nmt1-GFP-cnp1-lys1^+^ sad1-mCherry::kan^R^ leu1-32 ufd1-73* | This study |
| **YKK1124** | *h^-^ lys1::nmt1-GFP-cnp1-lys1^+^ sad1-mCherry::kan^R^ leu1-32 cdc48-353* | This study |
| **YKK1129** | *h^-^ GFP-cnp1::hph^R^ sad1-mCherry::kan^R^ leu1-32* | This study |
| **YKK1131** | *h^-^ GFP-cnp1::hph^R^ sad1-mCherry::kan^R^ leu1-32 ufd1-73* | This study |
| **YKK1125** | *h^-^ GFP-cnp1::hph^R^ sad1-mCherry::kan^R^ leu1-32 cdc48-353* | This study |
| **YKK1269** | *h^-^ cnt1/TM(NcoI)-ura4^+^ ura4-DS/E GFP-cnp1::hph^R^ leu1-32* | This study |
| **YKK1250** | *h^-^ cnt1/TM(NcoI)-ura4^+^ ura4-DS/E GFP-cnp1::hph^R^ leu1-32 urd1-73* | This study |
| **YKK1252** | *h^-^ cnt1/TM(NcoI)-ura4^+^ ura4-DS/E GFP-cnp1::hph leu1-32 cdc48-353* | This study |
| **YKK1241** | *h^-^ cdc48-GFP::kan^R^ ura4-D18 leu1-32 [pAU-ufd1-RFP]* | This study |
| **YKK1242** | *h^-^ GFP-cnp1::hph^R^ ura4-D18 leu1-32 [pAU-ufd1-RFP]* | This study |
| **AW3** | *h^-^ ufd1-GFP-ura4^+^ ura4-D18 leu1-32* | This study |
| **YKK1196** | *h^-^ ufd1-GFP-ura4^+^ ura4-D18 leu1-32 cdc48-353* | This study |
| **FY17243** | *h^-^ cdc48-GFP::kan^R^ leu1-32* | YGRC/NBRP |
| **YKK1197** | *h^-^ cdc48-GFP::kan^R^ leu1-32 ufd1-73* | This study |
| **YKK1430** | *h^-^ lys1::nmt1-GFP-cnp1-lys1^+^ swi6-tdTomato::hph^R^ leu1-32 ufd1-73 ade6-M210* | This study |
| **YKK1158** | *h^-^ cnp3-tdTomato::kan^R^ ura4-D18 leu1-32 [pREP81-cnp3c-GFP-ufd1]* | This study |
| **YKK1415** | *h^-^ cdc48-GFP::kan^R^ cnp3-tdTomato::kan^R^ ura4-D18 leu1-32 [pREP81-cnp3c-GFP-ufd1]* | This study |
| **NK91** | *h+ ufd1-GFP-ura4^+^ ch16-imr3L<<tetO-ura4^+^ ade6-M210 leu1-32 ura4-D18 [pREP41-npl4-mCherry]* | This study |
| **YKK1511** | *h^-^ leu1-32 [pREP81-cnp3c-GFP-ufd1/pREP41-npl4-mCherry]* | This study |
| **YKK1512** | *h^-^ leu1-32 [pREP81-cnp3c-GFP-ufd1/pREP41-mCherry]* | This study |
| **NK5** | *h^-^ GFP-cnp1::hph ch16-imr3L<<tetO-ura4^+^ ade6-M210 leu1-32 ura4-D18 [pREP41-tetR-mCherry]* | This study |
| **NK6** | *h^-^ GFP-cnp1::hph^R^ ch16-imr3L<<tetO-ura4^+^ ade6-M210 leu1-32 ura4-D18 [pREP41-tetR-npl4-mCherry]* | This study |
| **NK19** | *h^-^ cdc48-GFP::kan^R^ ch16-imr3L<<tetO-ura4^+^ ade6-M210 leu1-32 ura4-D18 [pREP41-tetR-mCherry]* | This study |
| **NK20** | *h^-^ cdc48-GFP::kan^R^ ch16-imr3L<<tetO-ura4^+^ ade6-M210 leu1-32 ura4-D18 [pREP41-tetR-npl4-mCherry]* | This study |
| **AW48** | *h^-^ ufd1-73-GFP-ura4^+^ ura4-D18 leu1-32* | This study |
| **MS548** | *h^-^ GFP-cnp1::hph^R^ cnp3-mCherry-leu1+ leu1-32* | Suma *et al*., (2018) |
| **YKK1477** | *h^-^ GFP-cnp1::hph^R^ cnp3-mCherry-leu1+ leu1-32 ufd1-73* | This study |
| **YKK1483** | *h^-^ GFP-cnp1::hph^R^ cnp3-mCherry-leu1+ leu1-32 cdc48-353* | This study |

**Supplementary Table 2. PCR primer for Chip assay and plasmids used in this study**

| *cnt1 forward* | Kitagawa et al., 2014 |
| --- | --- |
| *cnt1 reverse* | Kitagawa et al., 2014 |
| *imr1 forward* | Takayama et al., 2008 (28) |
| *imr1 reverse* | Takayama et al., 2008 |
| *dg1 forward* | Kitagawa et al., 2014 |
| *dg1 reverse* | Kitagawa et al., 2014 |
| 18S *forward* | Kitagawa et al., 2014 |
| 18S *reverse* | Kitagawa et al., 2014 |
| *act1 forward* | Takayama et al., 2008 |
| *act1 reverse* | Takayama et al., 2008 |
| *ura4 forward* | 5’-TACCTTTGGGACGTGGTCTC-3’ |
| *ura4 reverse* | 5’-CCCGTCTCCTTTAACATCCA-3’ |
| *subtelomere forward* | Hayashi et al., 2009 (29) |
| *subtelomere reverse* | Hayashi et al., 2009 |
| pYKK-1 | pREP81-cnp3c-GFP-ufd1 |
| pYKK-3 | pREP41-cnp3c-ufd1 |
| pYKK-5 | pREP41-ufd1 |
| pYKK-6 | pREP41-npl4-mCherry |
| pYKK-7 | pAU-ufd1-RFP |
| pYKK-8 | pREP41-tetR-mCherry |
| pYKK-9 | pREP41-tetR-ufd1-mCherry |

**Supplementary Legends**

**Supplementary Fig. 1. *ufd1-73* mutants are sensitive to over-expression of CENP-A/Cnp1.**

(A) Effect of over-expression of Cnp1 and histone H3. Strains ectopically expressing Cnp1 or histone H3 from the nmt1 promoter were plated on EMM plates with thiamine for repression (OFF) or without thiamine for derepression (ON) at 26 ℃ or 32 ℃. (B) Schematic illustration of the structure of Ufd1 protein and the position of the *ufd1-73* mutation. Partial amino-acid sequences of Ufd1 from five species are aligned. The mutation site in fission yeast Ufd1 is a single point mutation changing GGT (glycine) of the 55^th^ codon to GAT (aspartic acid). UFD1: ufd1 domain; SHP: p97 N domain binding and SIM: SUMO-interaction motif. (C) Localization of over-expressing GFP-Cnp1. Strains were grown to mid-log phase in liquid EMM+thiamine medium (repressed condition) at 26 ℃ and then transferred to EMM-thiamine medium (expressed condition) for induction of GFP-Cnp1 in the absence of thiamine. After the induction of GFP-Cnp1 for 18 hr, they were shifted to 32 ℃ for 8 hr. Sad1-mCherry: SPB maker. scale bar: 10 μm. (C-E) were observed under the same conditions. (D) The statistic analysis of (C). (E) The protein level of over-expressing GFP-Cnp1 by western blot. (F) Localization of Swi6 in *ufd1-73* under over-expressing Cnp1. Strains were grown as (C). scale bar: 2 μm. (G) Localization of over-expressing GFP-Cnp1 by ChIP analysis. Strains were grown as (C). The results are shown as a ratio to the EMM+thiamine (repressed condition) samples. All data represent the mean±s.e.m. (n = 3). *P*-values (unpaired *t*-test) comparing controls (WT): ***p* < 0.01.
