## Supplementary figures and images for "Cdc48 and its co-factor Ufd1 segregate CENP-A from centromeric chromatin and can induce chromosome elimination in the fission yeast *Schizosaccharomyces pombe*"

### supplement Figure (Ufd1)_YN0616_2023.pdf

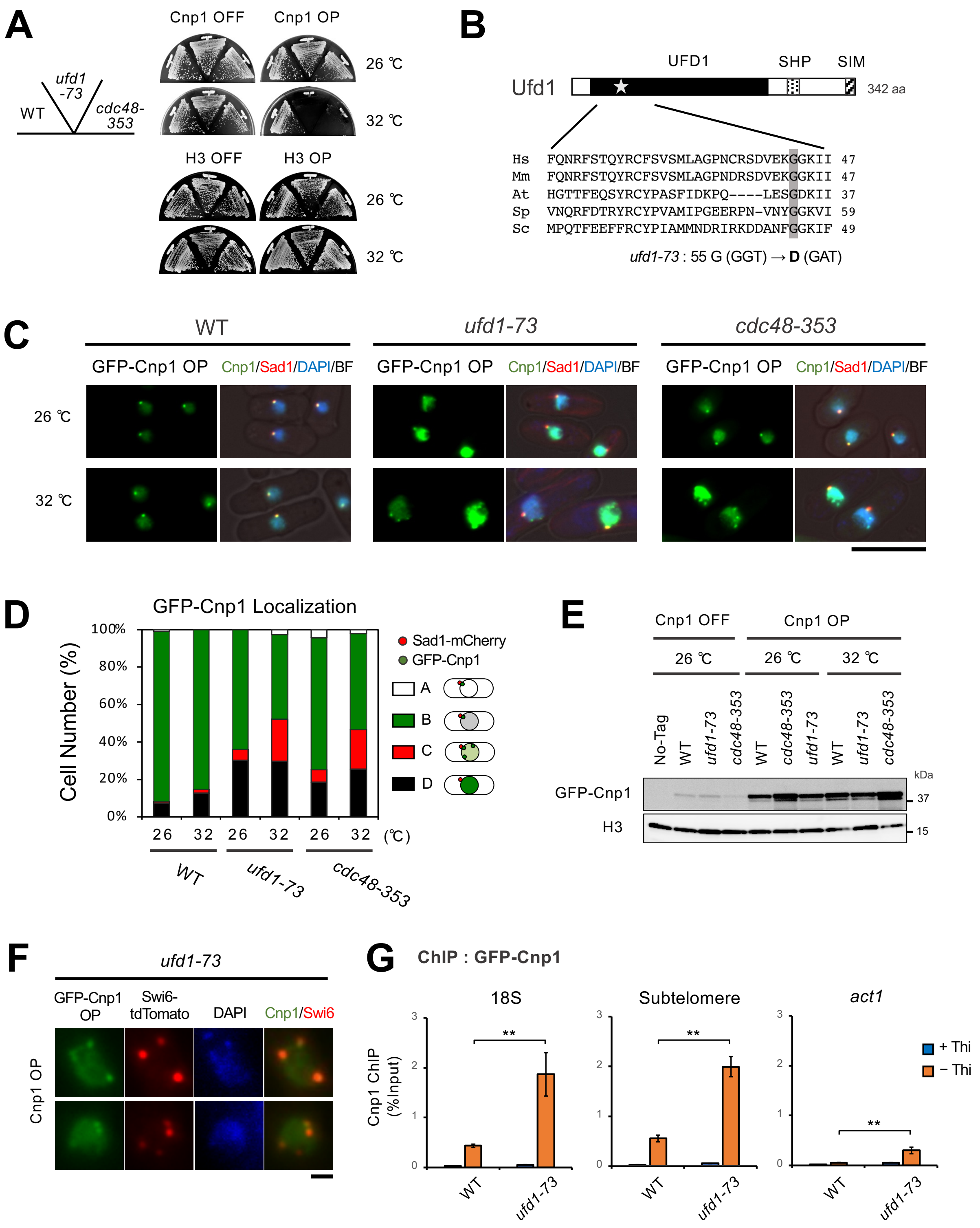
